# Tracking Human HSCs and MPPs in Mice Reveals Distinct Clonal Dynamics and Responses to Pre-transplant Conditioning

**DOI:** 10.64898/2026.02.10.705085

**Authors:** Mary Vergel Snodgrass, Jiya Eerdeng, Patrick Condie, Rong Lu

## Abstract

Blood and immune cell regeneration is sustained by hematopoietic stem and progenitor cells (HSPCs), which form the therapeutic basis of bone marrow transplantation. While the functional hierarchy of mouse HSPC subsets is well characterized, the distinct roles of human HSPC populations remain less well defined, particularly at clonal resolution and in the context of transplantation conditioning. While clonal tracking in humans and non-human primates has significantly advanced our understanding of hematopoietic dynamics, prior studies predominantly focused on CD34^+^ cells, a heterogeneous population of HSPCs. Moreover, secondary transplantation is considered the gold standard for distinguishing hematopoietic stem cells (HSCs) from multipotent progenitors (MPPs) in mice, but it has not been effectively utilized to study human HSPC populations.

To address this knowledge gap, we performed quantitative clonal tracking of purified human HSCs (hHSCs) and human MPPs (hMPPs) in NSGW41 mice across primary and secondary transplantation under no conditioning, busulfan, and irradiation. Consistent with prior studies, both hHSCs and hMPPs sustained long-term multilineage reconstitution and differed in engraftment rates. Our quantitative clonal analysis further revealed that hHSC clones generated more blood cells, initiated lymphoid production earlier, and exhibited more robust multilineage differentiation than hMPP clones. hHSC clones were also less sensitive to conditioning, maintaining stable lineage biases. Notably, busulfan and irradiation differentially affected the magnitude, lineage bias, and timing of hematopoietic reconstitution without altering engraftment. During secondary transplantation, hHSCs and hMPPs contributed comparably to hematopoietic reconstitution, but their overall output, particularly monocytes and T cells, was substantially reduced. In contrast to primary recipients, human chimerism of secondary recipients in the peripheral blood was diminished relative to the bone marrow and spleen, and more hHSPC clones contributed to hematopoiesis. Extramedullary hematopoiesis was observed in all secondary recipients, with comparable contributions from hHSC and hMPP clones.

Overall, this study provides insights into the distinct functions of hHSCs and hMPPs, the influence of conditioning, and the inefficiency of human hematopoiesis through serial transplantation. These findings advance our understanding of human hematopoiesis and provide a framework for utilizing and optimizing experimental models, improving transplantation conditioning strategies, and informing the preclinical evaluation of HSC-based cell and gene therapies.

## Background

Human hematopoietic stem and progenitor cells (hHSPCs) sustain lifelong blood and immune cell production and are central to bone marrow (BM) transplantation therapies ^1–5^. Within the hHSPC compartment, human hematopoietic stem cells (hHSCs) are defined by their long-term self-renewal and multilineage differentiation potential ^6,7^. They give rise to human multipotent progenitors (hMPPs) that also exhibit robust multilineage differentiation potential but limited engraftment capacity ^7–10^. While the functional roles and distinctions of mouse HSCs and MPPs are well characterized, the dynamic contributions of hHSCs and hMPPs to blood cell production remain less well defined ^11–15^. Previous studies in humans, non-human primates, and human xenograft models primarily used CD34⁺ or lineage-depleted cells, which comprise a mixture of hHSCs, hMPPs, and many types of downstream progenitors ^16–18^. In addition, human studies are largely limited to individuals with underlying diseases ^13,19^, studies in non-human primates are constrained by small sample sizes ^20–22^, and findings from mouse HSPCs may not be directly translatable to human HSPCs ^14,23^.

Xenograft mouse models have been widely used as preclinical platforms to study human hematopoiesis in vivo ^24–26^. These models allow for evaluating the lineage potential, self-renewal capacity, and differentiation dynamics of hHSPCs across multiple tissues and time points. Secondary transplantation, where HSPCs are recovered from primary recipients and transplanted into secondary recipients, provides a rigorous assay for assessing long-term self-renewal ^27^. This approach is routinely used to distinguish HSCs from MPPs in mouse models, but has not been systematically applied to the study of human HSC and MPP clones. In addition, it remains unclear how hHSPCs contribute to extramedullary hematopoiesis (EMH), the production of blood cells outside the BM ^28,29^. EMH occurs in both mice and humans in response to hematopoietic stress or disease, and most commonly manifests in the spleen ^30,31^. However, few studies have examined the involvement of additional anatomical sites in EMH or evaluated the contributions of individual hHSC and hMPP clones to this process.

A critical variable in both experimental and clinical transplantation is thepre-transplant conditioning, such as busulfan or irradiation, which is used to eradicate recipient HSPCs to facilitate donor cell engraftment ^32–36^. While these regimens are known to modulate the host environment and influence donor cell functions in mice ^12,37,38^, their effects on hHSPCs have not been directly investigated in a controlled experimental setting, particularly at the clonal level. Recent advances in lentiviral barcoding and clonal tracking have enabled lineage tracing of individual hHSPC clones following transplantation ^39–42^. These tools provide longitudinal, quantitative, and high-resolution analysis of how hHSPCs respond to conditioning regimens and contribute to hematopoiesis over time in both canonical and extramedullary niches.

In this study, we performed clonal tracking of purified hHSCs and hMPPs following primary and secondary transplantation into NSGW41 mice either unconditioned, or treated with busulfan or irradiation. By analyzing clonal contributions across peripheral blood, BM, spleen, and extramedullary sites, we identified distinct roles of hHSCs and hMPPs in hematopoietic reconstitution, revealed how pre-transplant conditioning influences their hematopoietic reconstitution, and delineated the strengths and limitations of xenograft mouse models for studying human hematopoiesis.

## Methods

### Isolation of hHSCs and hMPPs

Umbilical cord blood was obtained from the University of California Los Angeles (UCLA) Blood and Platelet Center. Mononuclear cells were isolated by Ficoll® Paque Plus (GE Healthcare/Cytiva) density gradient centrifugation. Isolated cells were then enriched in human hematopoietic progenitors by removing lineage-committed blood cells using the EasySep® Human Progenitor Cell Enrichment Kit with Platelet Depletion (StemCell Technologies) on the RoboSep®, following the manufacturer’s instructions. Human hematopoietic progenitors-enriched cells were subsequently sorted into human hematopoietic stem cells (hHSCs) and human multipotent progenitor cells (hMPPs) using fluorescence-activated cell sorting (FACS) with the following antibodies: hCD34-APC (clone 581), hCD38-APC-Cy7 (clone HB-7), hCD45RA-BV737 (clone HI100), and hCD90-PE-Texas Red (clone 5E10) (Figure S1A). All antibodies were purchased from BioLegend. Flow cytometry analysis confirmed that the purity of the sorted hHSCs and hMPPs exceeded 99%.

The cell surface marker combinations for hHSPCs:

hHSCs: hCD34^+^ / hCD38^-^ / hCD45RA^-^ / hCD90^+^

hMPPs: hCD34^+^ / hCD38^-^ / hCD45RA^-^ / hCD90^-^

### Lentiviral Transduction of Clonal Tracking Barcodes

The design and application of the barcode library have been described in detail previously ^39,42–45^. Prior to lentiviral transduction, sorted hHSCs and hMPP were plated in separate wells. Cells were then stimulated ex vivo for 24 hours in StemSpan™ SFEM (StemCell Technologies) supplemented with 10X GlutaMAX™ (Thermo Fisher Scientific) and 10 U/mL penicillin-streptomycin. The stimulated cells were then transduced in a 96-well U-bottom plate containing the stimulation media supplemented with 100 ng/mL IL-6 (Thermo Fisher Scientific) and 100 ng/mL StemSpan™ CC110 Cytokine Cocktail (StemCell Technologies). Each well was transduced with lentivirus carrying a unique barcode library. After 16 hours of transduction, cells were washed three times and resuspended in PBS containing 2% FBS.

### Xenotransplantation of hHSPCs

All animal procedures were conducted in accordance with protocols approved by the Institutional Animal Care and Use Committee of the University of Southern California (USC). NSGW41 mice (NOD.Cg-*Kit^W-41J^ Prkdc^scid^ Il2rg^tm1Wjl^*/WaskJ, JAX stock #026497) were purchased from the Jackson Laboratory. Mice were housed under sterile conditions and provided with Uniprim food (Envigo) and acidified water at the time of transplantation.

Adult mice (6–12 weeks old) were conditioned either by total body irradiation (100 cGy) or intraperitoneal injection of busulfan (20 mg/kg; Sigma-Aldrich) administered 24 hours prior to transplantation. Each primary recipient mouse received a retro-orbital injection of 7,000 sorted and barcoded hHSCs and 18,000 sorted and barcoded hMPPs in 100 uL of PBS with 2%FBS. Five mice were transplanted per group, including an unconditioned group (Supplementary Table 1).

Five months after primary transplantation, BM was harvested, and hHSCs and hMPPs were then isolated, sorted and transplanted into multiple secondary recipients. Twelve months after secondary transplantation, BM, spleen, peripheral blood, and samples of any tissue exhibiting prominent extramedullary hematopoiesis were collected for further analysis. For secondary recipients, data were obtained from 17 busulfan-conditioned, 11 unconditioned, and 6 irradiated mice that exhibited donor engraftment (Supplementary Table 2).

### Peripheral Blood Cell Analysis

Blood samples were collected via tail vein bleeding into PBS containing 10 mM EDTA. To deplete red blood cells, 2% dextran was added to sediment erythrocytes, and the remaining leukocyte-enriched fraction was treated with ammonium-chloride-potassium (ACK) lysis buffer on ice for 5 minutes to remove residual erythrocytes. The following antibodies were used for flow cytometric analysis of blood cells: mCD45-A700 (clone A20, BioLegend), hCD45-APC780 (clone HI30, Thermo Fisher Scientific), hCD3-PerCP-Cy7 (clone UCHT1, Thermo Fisher Scientific), hCD8-PE-Cy5 (clone HIT8a, BioLegend), hCD4-PE-Dazzle (clone OKT4, BioLegend), hCD19-PE (clone SJ25C1, BioLegend), hCD13-APC (clone MHCD1305, Thermo Fisher Scientific), hCD33-APC (clone P67.6, BioLegend), and hCD66b-PerCP-Cy5.5 (clone G10F5, BioLegend). After one hour of incubation, cells were resuspended in PBS containing 2% FBS (VWR) supplemented with DAPI (4′,6-diamidino-2-phenylindole, 1:10,000), then analyzed and sorted using a FACSAria™ II cell sorter.

The cell surface marker combinations for human hematopoietic cells:

Granulocytes: mCD45^-^ / hCD45^+^ / hCD3^-^ / hCD19^-^ / hCD66^+^

Monocytes: mCD45^-^ / hCD45^+^ / hCD3^-^ / hCD19^-^ / hCD66^-^ / hCD13^+^ / hCD33^+^

B cells: mCD45^-^ / hCD45^+^ / hCD3^-^ / hCD19^+^

T cells: mCD45^-^ / hCD45^+^ / hCD3^+^ / hCD19^-^

### Cell Isolation from Solid Tissues

Mononuclear cells (MNCs) were isolated from the spleen, bone marrow, and extramedullary hematopoietic (EMH) sites of recipient mice. Fresh spleens were mechanically dissociated in PBS supplemented with 2% FBS and passed through a 70 µm cell strainer to obtain single-cell suspensions. Cells were then stained with anti-hCD45 and anti-mCD45 antibodies. For non-splenic EMH sites, small tissue fragments exhibiting visible areas of cell growth or infiltration were collected and dissociated into single-cell suspensions either by mechanical dissociation or enzymatic digestion using Collagenase/Dispase (1 mg/mL). Dissociated cells were then stained with the anti-hCD45 and anti-mCD45 antibodies. Human MNCs were subsequently isolated by fluorescence-activated cell sorting (FACS).

### Barcode Extraction and Analysis

Genomic DNA was extracted from sorted cells and amplified using Phusion™ PCR Master Mix (Thermo Fisher Scientific) as previously described ^15,39,42^. PCR products were purified using SPRI magnetic beads (Beckman Coulter) following the manufacturer’s instructions. The purified amplicons were then sequenced at the Children’s Hospital Los Angeles Sequencing Core Facility using Illumina NextSeq kit. Sequencing data were processed as previously described ^39,42^. Briefly, custom Python scripts were used to process FASTQ files and remove low-quality reads. Reads corresponding to each sample were identified based on primer indices. Barcode sequences were then identified by exact matching to the expected library ID.

Sequencing data were integrated with flow cytometry data to quantify the abundance of each clone, as described below. Unique DNA barcodes with abundances greater than 0.001% in any cell type were included in the downstream analysis.

Clonal abundance of blood cells (Figure 2-4) was calculated relative to all mononuclear cells (MNCs) in the peripheral blood of each mouse using the following formula:

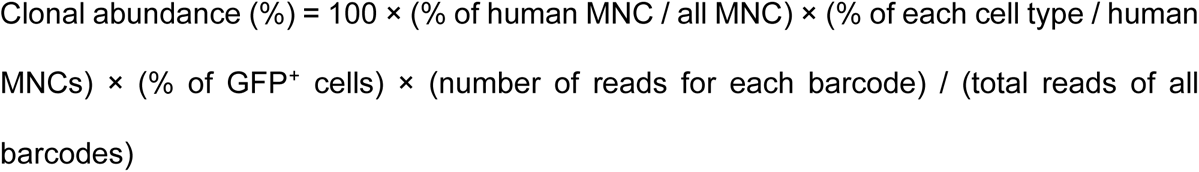

Clonal abundance comparing across tissues (Figure 5F-5G) was calculated relative to all human mononuclear cells (MNCs) in each tissue using the following formula:

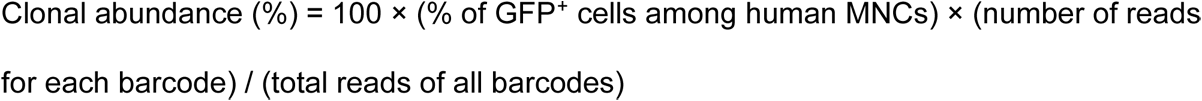

### Statistical Analysis

Flow cytometry data were analyzed using FlowJo (version 10.9; BD Biosciences) and GraphPad Prism (version 10.4 for Windows; GraphPad Software). Clonal data analysis was conducted using custom Python scripts, with visualizations generated using the ‘matplotlib’ and ‘seaborn’ libraries. Data are presented as the mean + standard error of the mean (SEM). Statistical significance was defined as p < 0.05. The statistical methods used are indicated in the corresponding figure legends.

## Results

### Blood Cell Production of hHSPCs is Influenced by Pre-transplant Conditioning

To investigate how conditioning regimens affect blood cell production of hHSPCs, we isolated hHSCs and hMPPs from umbilical cord blood ^8,46^ and labeled them with distinct lentiviral barcoding libraries, which were distinguishable by barcode sequencing but not by flow cytometric analysis, as all libraries expressed GFP ^39,42^ (Figures 1A and S1A). 7,000 hHSCs and 18,000 hMPPs were then pooled and transplanted as a mixed hHSPC population into each NSGW41 recipient mouse, either unconditioned or conditioned with irradiation or busulfan prior to transplantation (Figure 1A, Supplementary Table 1). Approximately 20% of hHSCs and hMPPs were barcoded, and their contribution to hematopoiesis was comparable to that of non-barcoded cells (Figure S2).

**Figure 1.**
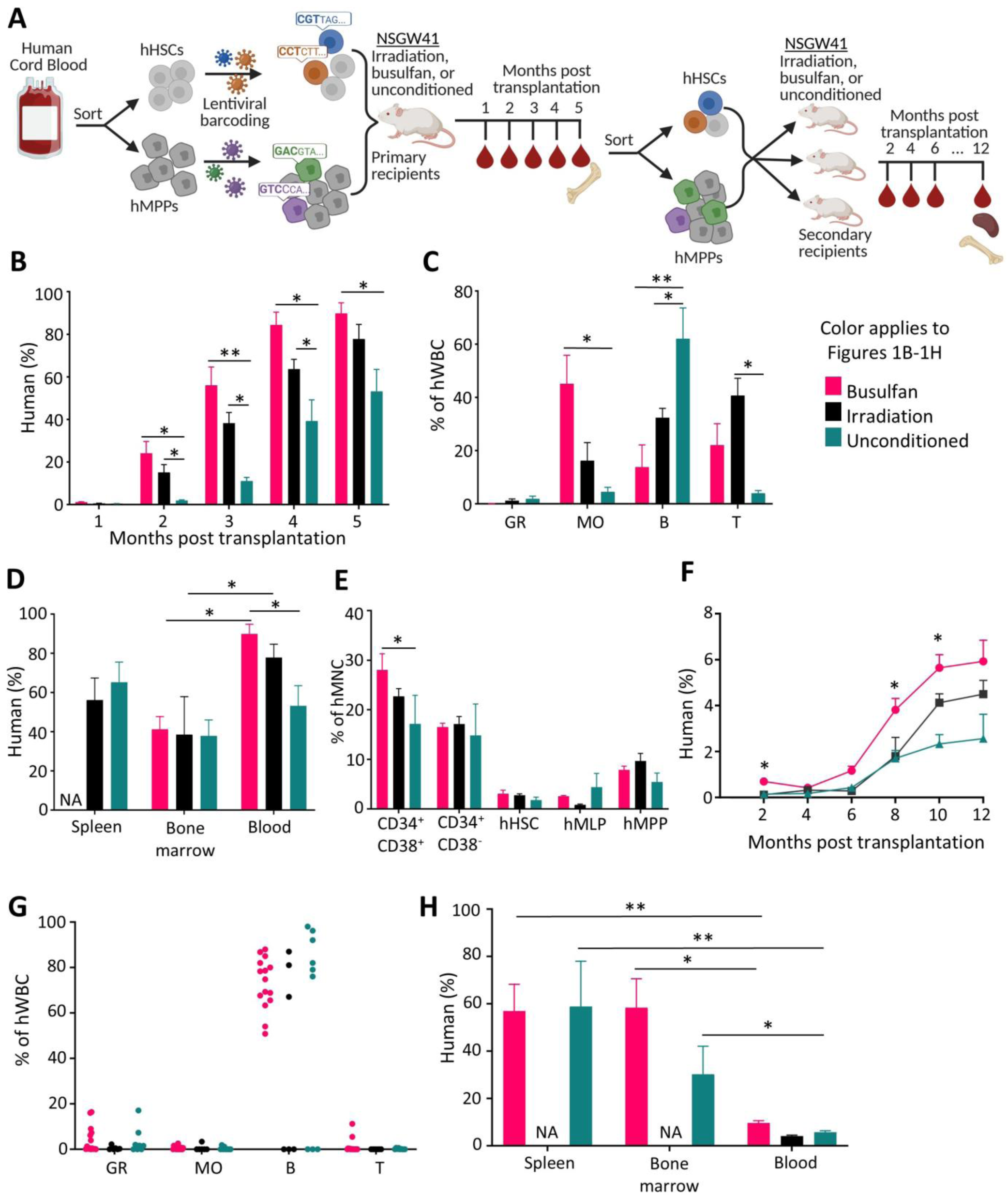
Blood cell production by hHSPCs is influenced by pre-transplant conditioning. **(A)** Schematic overview of the experimental workflow. For primary transplantation, five mice were transplanted per conditioning group. For secondary transplantation, data were collected from 17 busulfan-conditioned, 11 unconditioned, and 6 irradiated recipients. **(B)** Percentage of human hematopoietic cells (hCD45^+^) in the peripheral blood of primary recipient mice over time. **(C)** Composition of human-derived hematopoietic cells in the peripheral blood of primary recipients at 5 months post-transplantation. **(D)** Percentage of human hematopoietic cells in various tissues of the primary recipients at 5 months post-transplantation. **(E)** Composition of human-derived hematopoietic progenitor cells in the BM of the primary recipients at 5 months post-transplantation. **(F)** Percentage of human hematopoietic cells in the peripheral blood of secondary recipient mice. Significant differences were observed between the busulfan and unconditioned groups at the three indicated time points. **(G)** Composition of human-derived hematopoietic cells in the peripheral blood of secondary recipients at 12 months post-transplantation. **(H)** Percentage of human hematopoietic cells in various tissues of the secondary recipients at 12 months post-transplantation. **(B - F**, **H)** show mean + SEM. Differences among the three conditioning regimens were analyzed by either repeated-measures (**B, F**) or not repeated (**C - E, H**) two-way ANOVA with Tukey post-hoc tests; *p<0.05, **p<0.01. Abbreviations: NA, not available; hWBC, human white blood cells; hMNC, human mononuclear cells; hMLP, human multi-lymphoid progenitors; GR, granulocytes; MO, monocytes; B, B cells; T, T cells.

Monthly peripheral blood analyses revealed that pre-transplant conditioning significantly increased the abundance of hHSPC-derived cells, with the highest levels observed in busulfan-treated recipients (Figure 1B and S1B). Furthermore, the conditioning regimen influenced lineage bias of hHSPCs, with busulfan skewing differentiation toward monocytes, irradiation promoting T cell production, while the absence of conditioning favored B cell lymphopoiesis (Figures 1C and S1C). Despite these differences in peripheral blood cell output, total human chimerism in the BM and spleen was surprisingly not influenced by the conditioning regimen (Figure 1D). Further analysis revealed that hHSC and hMPP reconstitution in the recipient mouse BM was also unaffected by the conditioning regimen (Figure 1E). However, busulfan-treated mice exhibited a significantly higher frequency of CD38⁺CD34⁺ progenitor cells in the BM compared to unconditioned mice (Figure 1E), consistent with their increased human chimerism and myeloid output in the peripheral blood (Figures 1B and 1C). Collectively, these findings suggest that pre-transplant conditioning modulates the differentiation, but not engraftment, of hHSPCs, influencing both the quantity and lineage bias of their blood cell production.

### Reduced Blood Cell Production in Secondary Recipients

Five months after the primary transplantation, hHSCs and hMPPs were collected from each primary recipient and transplanted into multiple secondary recipient mice that had received the same pre-transplant conditioning as the corresponding primary recipients (Figure 1A, Supplementary Table 2). Human chimerism in the peripheral blood of secondary recipients was low initially and increased after six months post-transplantation (Figure 1F), a marked delay compared to the primary recipients (Figure 1B). Consistent with primary transplantation, busulfan conditioning resulted in the highest levels of human chimerism in the secondary recipients, followed by irradiation, while unconditioned recipients exhibited the lowest chimerism (Figure 1F). Overall, human chimerism in the peripheral blood of the secondary recipients was below 6% across all experimental groups, substantially lower than the 50-80% chimerism observed in the primary recipients (Figures 1B and 1F).

In addition to chimerism, the types of blood cells produced by hHSPCs in the secondary recipients were also strikingly different from those in the primary recipients (Figures 1C and 1G). While human monocytes and T cells were robustly produced in primary recipients at 4 and 5 months post-transplantation (Figures 1C and S1C), these lineages were undetectable in most secondary recipients even up to 12 months post-secondary transplantation (Figure 1G). In contrast, granulocyte production was elevated in some busulfan-treated and unconditioned secondary recipients compared to the primary recipients (Figures 1C, S1C, and 1G). B cell production was abundant in both primary and secondary recipients. However, the impact of pre-transplant conditioning on B cell production was evident in primary recipients but not in secondary recipients (Figures 1C and 1G). Furthermore, in secondary recipients, human chimerism was substantially higher in the BM and spleen than in the peripheral blood (Figure 1H), opposite to what is observed in primary recipients (Figure 1D). These results indicate that the blood cell production capacity of hHSCs and hMPPs was substantially altered upon secondary transplantation, which was not reported by earlier xenograft studies using CD34⁺ or lineage-depleted cells ^47–49^.

### Long-Term Multi-Lineage Differentiation of hHSCs and hMPPs

While mouse MPPs contribute to hematopoiesis for less than four months post-transplantation ^1,50,51^, we observed multi-lineage blood cell contributions from hMPPs up to five months in primary recipients and up to twelve months in secondary recipients (Figure 2). Overall, the numbers of engrafted and differentiating hHSC and hMPP clones were low and comparable in primary and secondary recipients (Figure 2A and S3A). The low clone number is consistent with previous clonal tracking studies, considering the number of donor cells transplanted ^8,52,53^. Given that 7,000 hHSCs and 18,000 hMPPs were transplanted into each primary recipient and their similar transduction rate (Figure S2A), our data suggest a higher engraftment efficiency of hHSCs compared to hMPPs, consistent with previous findings ^8,52^.

**Figure 2.**
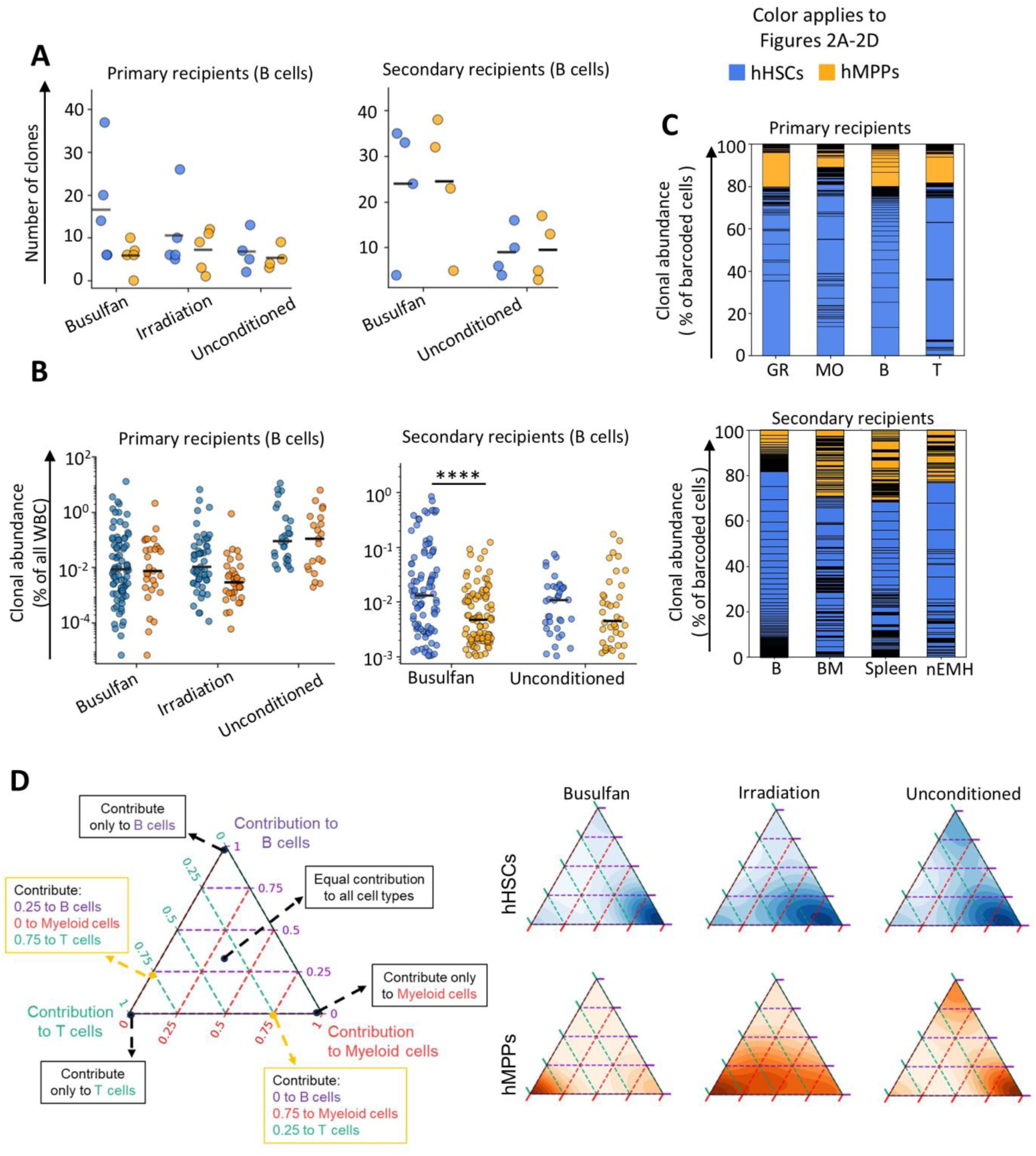
Long-term multi-lineage differentiation of hHSCs and hMPPs. Comparing clonal contributions of hHSCs and hMPPs to the primary recipients 5 months post-transplantation and to the secondary recipients 12 months after secondary transplantation. **(A)** Number of hHSC and hMPP clones contributing to B cells in the peripheral blood of primary and secondary recipients. Each dot represents data from one recipient mouse. Black bars indicate the means. **(B)** Clonal contributions of hHSCs and hMPPs to B cells in the peripheral blood of primary and secondary recipients. Each dot represents a unique clone. Black bars indicate the medians; Mann–Whitney test. **p<0.01. **(C)** Relative contributions of hHSC and hMPP clones to various hematopoietic cell types and tissues in the primary and secondary recipients. Each section within the columns represents one clone. Clones are ordered by their B cell contribution. **(D)** Lineage bias of hHSC and hMPP clones in primary recipients treated with busulfan, irradiation, or unconditioned. The density in the ternary plot reflects the number of clones. Abbreviations: hWBC, human white blood cells; hMNC, human mononuclear cells; GR, granulocytes; MO, monocytes; B, B cells; T, T cells; nEMH, non-splenic extramedullary hematopoietic sites; BM: bone marrow.

In addition to higher engraftment rate, hHSC clones produced similar or greater numbers of hematopoietic cells compared with hMPP clones in primary and secondary recipients (Figures 2B and S3B). Overall, hHSCs contributed more than hMPPs across all analyzed cell populations in both primary and secondary recipients (Figures 2C and S3C-3D). Furthermore, hMPP clones, but not hHSC clones, substantially altered lineage biases in response to conditioning (Figure 2D). In unconditioned mice, hHSC and hMPP clones displayed similar distributions of lineage bias (Figure 2D). However, after irradiation or busulfan conditioning, many hMPP clones altered their lineage biases, generating proportionally more T cells and fewer myeloid cells (Figure 2D). Such changes were not observed in hHSC clones, which maintained similar lineage bias distributions across unconditioned, irradiated, and busulfan-treated recipients (Figure 2D). Together, these results suggest that hMPP clones are more sensitive to transplantation conditioning than hHSC clones, while hMPP and hHSC clones exhibit similar lineage biases in the absence of conditioning.

Collectively, despite differences in clonal contribution and lineage bias, both hHSC and hMPP clones demonstrated the capacity for long-term, multi-lineage blood cell reconstitution (Figure 2), in contrast to their murine counterparts. This is consistent with previous population-level studies showing that hMPPs engraft less efficiently than hHSCs but support long-term, multi-lineage hematopoiesis ^8,52,54^. Yet, the long-term analysis results in mice may not be directly translatable to humans, given the substantially longer lifespan of humans compared with mice.

### Temporal Dynamics of Blood Cell Regeneration

As mouse MPPs initiate hematopoiesis earlier than mouse HSCs following transplantation ^50,51^, we next examined the temporal dynamics of blood cell production by hHSC and hMPP clones in primary recipients at months 3, 4, and 5 post-transplantation (Figure 3), focusing on monocytes and B cells, the most abundant myeloid and lymphoid cell types across all time points, respectively (Figures 1C and S1C). We clustered hHSC and hMPP clones based on temporal changes in clonal abundance and compared the dynamic profiles of clusters containing the highest numbers of clones. Our results suggest that hMPP clones were more likely to generate early monocytes than hHSC clones, whereas early B-cell output was more frequently initiated by hHSC clones (Figure 3A). Over time, more hHSC clones simultaneously contribute to B cells and monocytes compared with hMPP clones (Figures 3B and S4B).

**Figure 3.**
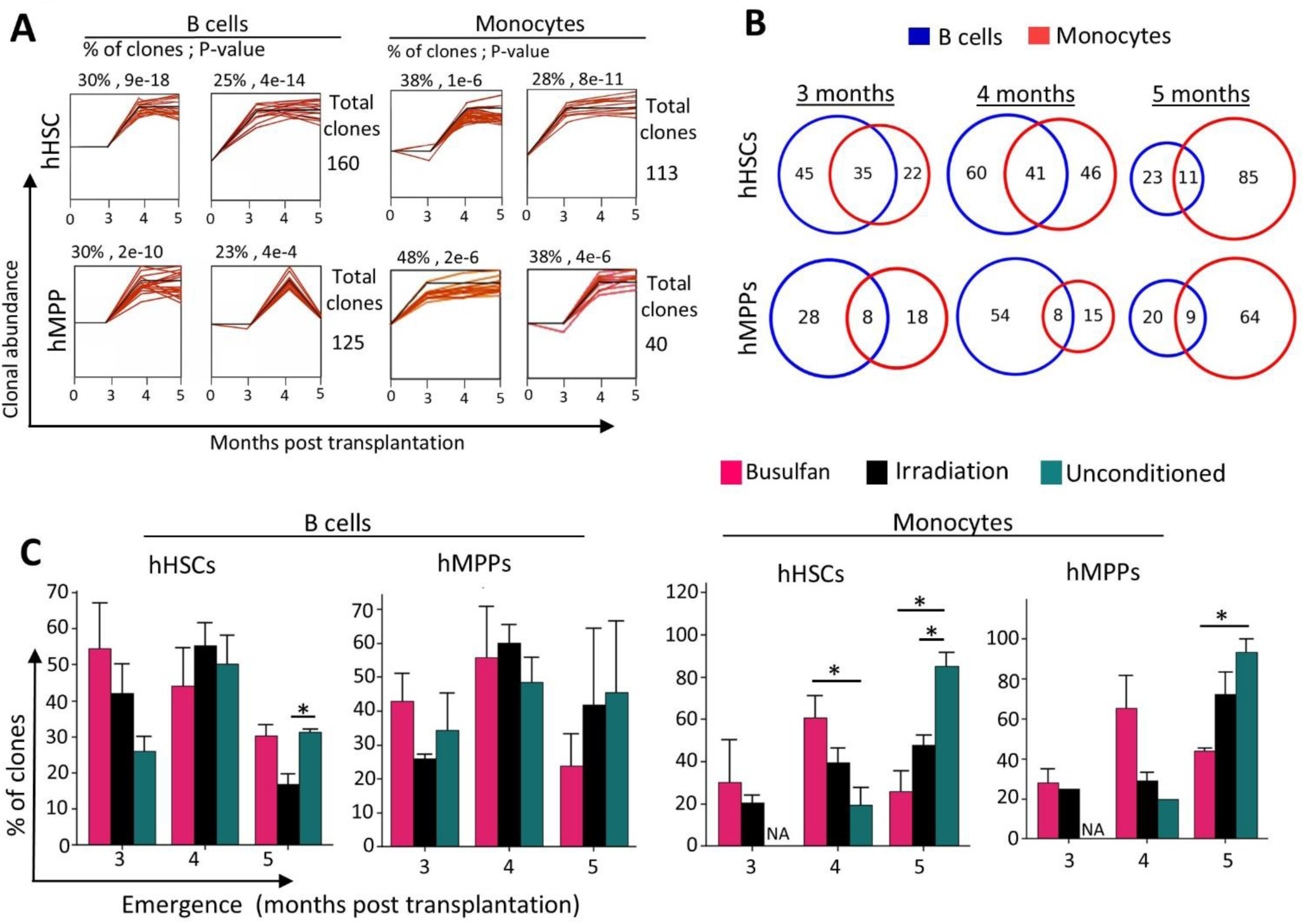
Temporal dynamics of hHSPC-derived blood cell production in primary recipients. **(A)** Short Time-series Expression Miner (STEM^55^) analysis showing temporal clustering of clonal contributions from hHSCs and hMPPs to B cells and granulocytes in peripheral blood. Each panel represents a cluster of clones with similar temporal dynamics. Shown are the two most prevalent temporal patterns within each group. Numbers above each panel denote the percentage of total clones in the cluster and the corresponding p-value relative to the expected proportion. Each red line represents a single hHSPC clone, and the black line shows the average temporal pattern for the cluster. Clones from all conditioned and unconditioned groups were included in this analysis. **(B)** Number of hHSC and hMPP clones that began contributing to B cells, monocytes or both at various time points post-transplantation. The percentage of overlap is shown in Figure S4B. **(C)** Percentage of hHSC and hMPP clones that began contributing to blood cells at various time points post-transplantation under different conditioning regimens. NA, not available, due to low cell numbers at that time point. Student’s t-test, ** p<0.01.

Our results also showed how pre-transplant conditioning regimens influence the temporal dynamics of blood cell regeneration by hHSPC clones. After irradiation- or busulfan-based conditioning, hHSPC clones exhibited similar temporal dynamics of contributing myeloid and lymphoid lineages with an initial increase followed by a plateau over time. (Figure S4A). In contrast, without conditioning, monocyte contributions from hHSPC clones continued to increase gradually even after B-cell production had peaked (Figure S4). Moreover, most hHSC and hMPP clones initiated monocyte production at 5 months post-transplantation in unconditioned recipients, which is significantly more than in conditioned recipients at the same time point (Figure 3C). These results suggest that engrafted hHSPCs exhibited delayed myelopoietic reconstitution in the absence of conditioning, highlighting the role of transplantation conditioning in promoting early myelopoiesis during hematopoietic recovery. The delayed human myelopoiesis, but not lymphopoiesis, in unconditioned mice may be related to the lack of endogenous lymphoid cells in this mouse model.

### Clonal Activation of hHSPCs upon Secondary Transplantation

In secondary recipients, we observed a substantial reduction in human chimerism in the peripheral blood, but not in the BM or spleen, suggesting reduced hematopoietic efficiency (Figures 1D and 1H). To investigate the underlying clonal dynamics during serial transplantation, we compared the clonal contributions to B cells in the peripheral blood between primary and secondary recipients, as B cells are the only blood cell type consistently and robustly produced by hHSPCs across both primary and secondary recipients (Figures 1C and S1C). Surprisingly, we found significantly more B cell contributing clones in secondary recipients than in the corresponding primary recipients (Figures 4A–4B, and S5A). Consistently, more hHSPC clones were detected in the spleens of secondary recipients (Figures 4A and S5A). In addition, clones detected exclusively in secondary recipients exhibited lower levels of B cell contribution compared to those found in both primary and secondary recipients (Figure 4C). Collectively, these results suggest that secondary transplantation activated previously quiescent or latent hHSPC clones, promoting their differentiation and clonal contribution.

**Figure 4.**
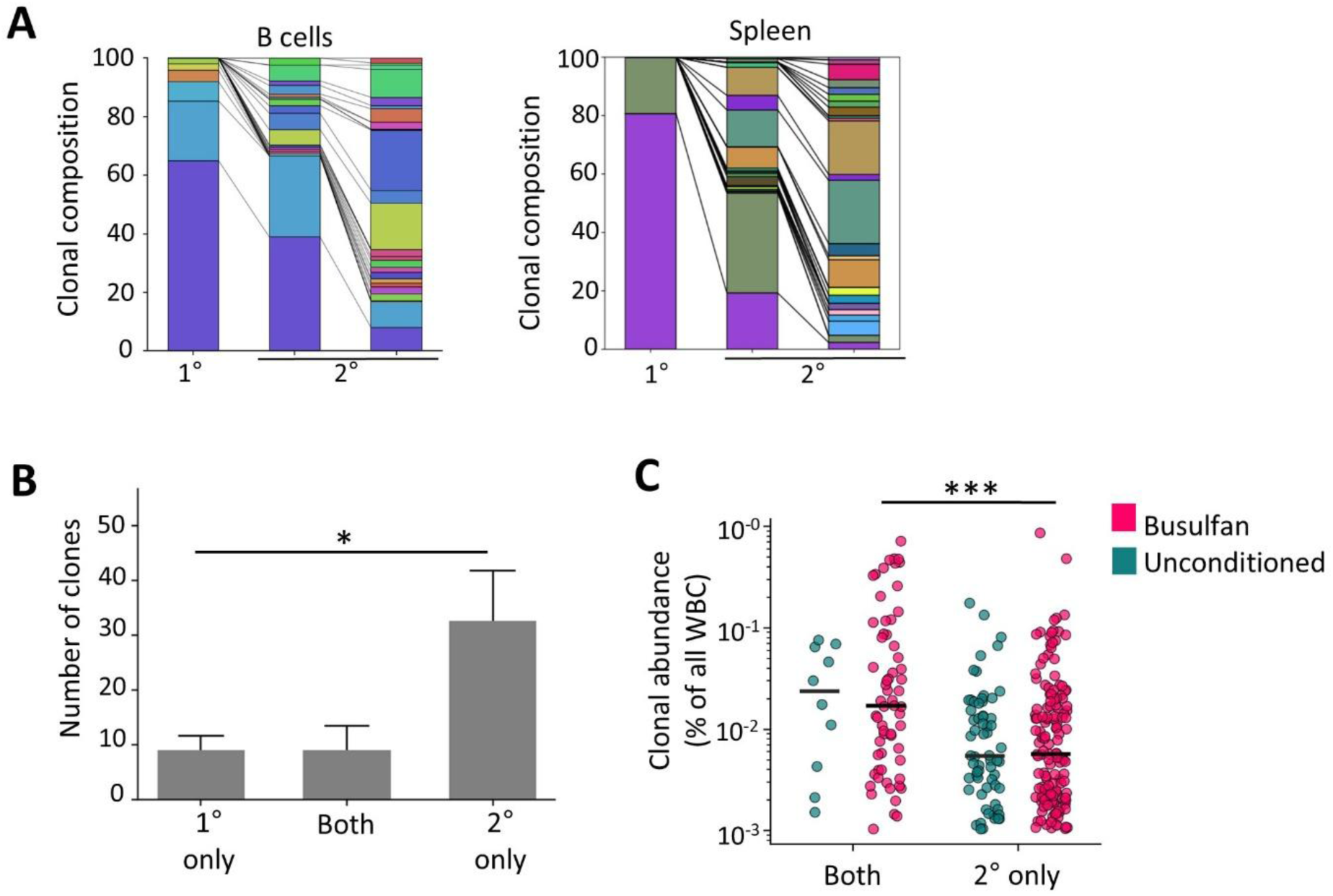
Hematopoietic contributions of hHSPC clones in primary and secondary recipients. Clones derived from hHSCs and hMPPs are combined in the analysis. **(A)** Clonal composition of human B cells in the peripheral blood (left) and human hematopoietic cells in the spleen (right), comparing a primary recipient (1°) with its derived secondary recipients (2°). Each color represents a unique clone. Each column shows one mouse. Additional mice are shown in Figure S5A. **(B)** Number of clones detected in peripheral blood B cells from primary (1°) and secondary (2°) recipients, showing those unique to each or shared between both. Clones from all conditioned and unconditioned groups were included in this analysis. Data are presented as mean + SEM; Student t-test; *p<0.05. **(C)** Clonal contribution of hHSPCs to peripheral blood B cells in the secondary recipients at 12 months post-transplantation, comparing clones that did or did not contribute to peripheral blood B cells in the primary recipients. Black bars indicate the means. *** p<0.001.

### Extramedullary Hematopoiesis Initiated by hHSPCs in secondary recipients

Five secondary recipient mice displayed substantial enlargement of the lymph nodes, ovaries, or lungs (Figure 5A), indicating EMH as observed in previous studies ^56–60^. Moreover, significantly enlarged spleens were observed in all secondary recipients (Figures 5B-5C), another indication of EMH ^60–62^. EMH can arise from insufficient hematopoiesis in the BM and has been reported in xenograft models ^29,31,63^, likely due to inadequate support for human hematopoiesis in the murine BM microenvironment ^29,64^. Our data now provide direct evidence of inefficient differentiation of hHSPCs in the BM of secondary recipient mice, as indicated by significantly reduced human chimerism in the peripheral blood compared to the BM and spleen (Figure 1H). In humans, EMH is associated with pathological conditions such as hematologic malignancies, BM failure, and systemic infections ^31,63^, highlighting its relevance as a compensatory hematopoietic response under stress and disease.

**Figure 5.**
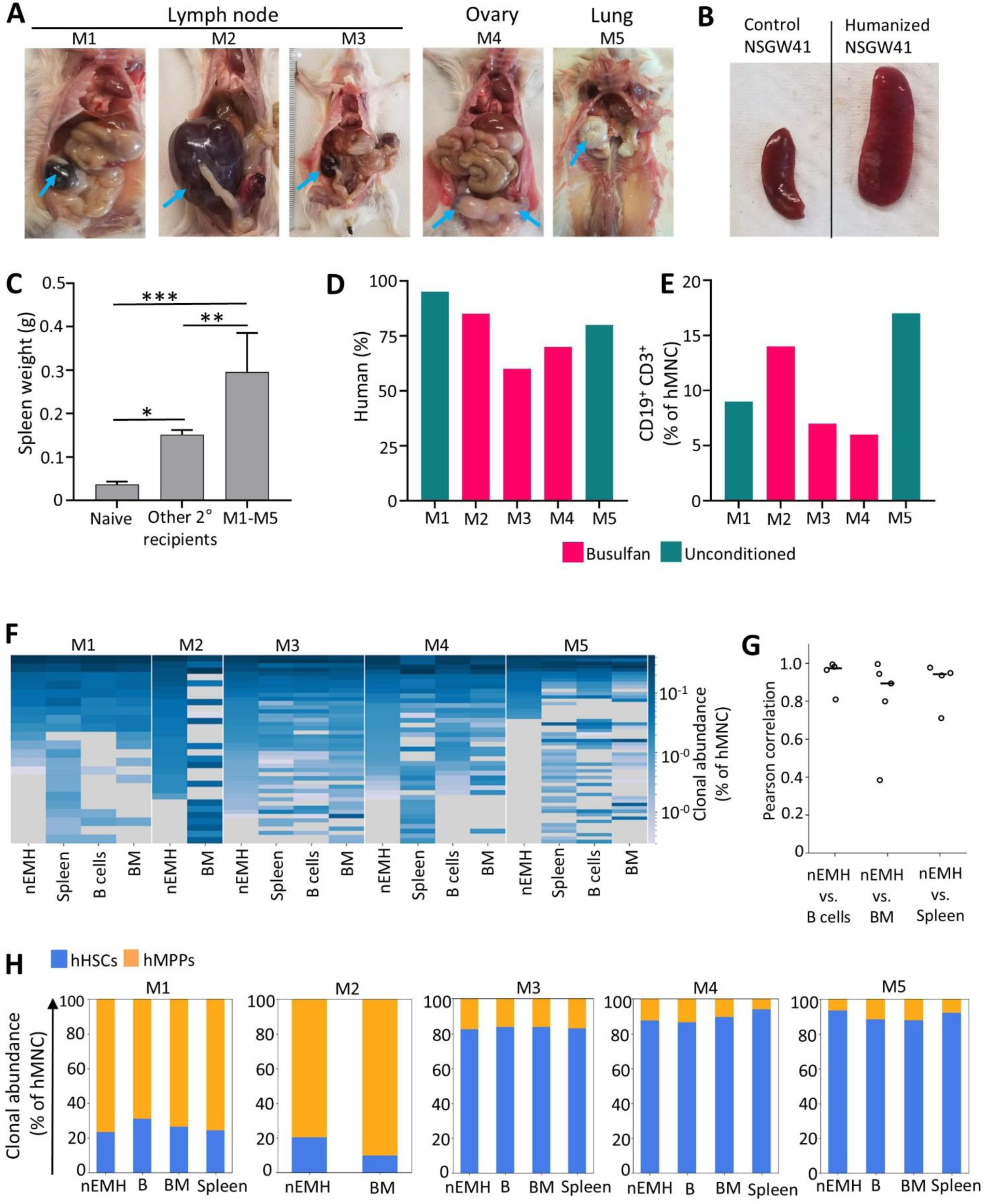
Extramedullary hematopoiesis (EMH) initiated by hHSPCs in the secondary recipients. Data were collected from five secondary recipients (M1–M5) exhibiting non-splenic extramedullary hematopoiesis (nEMH) at 12 months post-transplantation. **(A)** nEMH sites indicated by blue arrows. **(B)** Representative images of enlarged spleens from secondary recipient mice. **(C)** Spleen weights of secondary (2°) recipient mice with (n=5) and without (n=24) nEMH, compared with naïve control mice (n=4). Data are presented as mean + SEM; Student’s t-test; *p<0.05, **p<0.01, ***p<0.001. **(D)** Percentage of human hematopoietic cells (hCD45^+^) among all mononuclear cells (MNC) in nEMH sites of each mouse as highlighted in (A). Spleen data are shown in Figure S5B. **(E)** Frequency of CD19^+^CD3^+^ cells at the nEMH sites. **(F)** Clonal contribution across different tissues in mice with nEMH. Each row represents a unique clone. **(G)** Pearson correlation of hHSPC clonal composition between nEMH sites and hematopoietic tissues. Each dot represents an individual mouse. Black bars indicate the medians. **(H)** Relative contribution of hHSCs and hMPPs to nEMH sites and hematopoietic tissues. **(F-H)** B cells are derived from the peripheral blood. Abbreviations: B, B cells; nEMH, non-splenic extramedullary hematopoietic sites; BM: bone marrow.

To further characterize EMH, we examined the cellular composition at the EMH sites, including the spleens, lymph nodes, ovaries, and lungs. We found that approximately 75% of the cells in these sites were human hematopoietic cells (Figures 5D and S5B), in contrast to less than 6% human chimerism observed in the peripheral blood (Figure 1F). Notably, around 10% of the human cells at the non-splenic EMH sites co-expressed CD19 and CD3 (Figure 5E). This aberrant co-expression of both B cell (CD19) and T cell (CD3) markers was not detected in any other tissues, including the spleens of these mice, nor in any other xenografted mice. CD19^+^CD3^+^ cells have been associated with malignant phenotypes, including lymphoid tumors and leukemic blasts ^65–68^. Their presence at the non-splenic EMH sites suggests dysregulated lymphopoiesis in the extramedullary environment.

It has been speculated that EMH may be initiated by select subsets of hHSPC clones ^28^, yet this possibility lacks definitive experimental support. Our results demonstrate that the majority of hHSPC clones sustaining BM hematopoiesis also contributed to hematopoiesis at the EMH sites (Figure 5F). We observed a strong clonal correlation among extramedullary sites, BM, and peripheral blood, indicating a shared clonal composition (Figure 5G). In addition, hHSC and hMPP clones exhibited comparable contributions to extramedullary and BM hematopoiesis (Figure 5H), suggesting that both hHSC clones and hMPP clones are capable of migrating to EMH sites, adapting to the local microenvironment, and sustaining hematopoiesis outside the BM.

## Discussion

In this study, we performed clonal tracking of hHSCs and hMPPs in primary and secondary xenografted mice treated with different pre-transplant conditioning regimens. Our findings provide new insights into three key areas: the cellular dynamics of human hematopoietic reconstitution, the influence of pre-transplant conditioning, and the utility of xenograft mouse models for studying human hematopoiesis. In contrast to previous clonal tracking studies primarily using CD34⁺ or lineage-depleted cells^16–18^, this study specifically tracked purified hHSCs and hMPPs, enabling a direct and quantitative comparison of their clonal behavior and functional output. We show that hHSC and hMPP clones exhibit marked differences in blood cell contribution levels (Figures 2B-2C), temporal dynamics (Figure 3), and multi-lineage reconstitution (Figures 2D and 3B). In particular, hMPP clones, but not hHSC clones, substantially altered lineage biases in response to conditioning regimens (Figure 2D), consistent with previous studies indicating the HSCs are more resistant to environmental stress ^69,70^. Furthermore, the origin of EMH was poorly understood particularly in humanized mouse models ^31^. Our study demonstrates that both hHSC and hMPP clones exhibit consistent blood cell reconstitution between primary and secondary recipients (Figures 2A–2C) and contribute comparably to BM hematopoiesis and EMH (Figures 5F-5G), providing insights into the cellular origin of human EMH. Together, these findings advance our understanding of the distinct and shared functional capacities of hHSPC subpopulations and offer new perspectives on the cellular dynamics underlying hematopoietic reconstitution following BM transplantation.

This study also demonstrates how transplantation conditioning influences the clonal dynamics of hHSPCs. Conditioning regimens such as busulfan and irradiation are widely used in clinical transplantation to facilitate donor cell engraftment ^71–75^. Our data reveal that both busulfan and irradiation significantly and differentially alter the levels of human blood cell reconstitution (Figures 1B and 1F), influence lineage bias (Figures 1C and 2D), and shift the temporal dynamics of blood cell emergence (Figures 3C and S4). In particular, our results demonstrate that pre-transplant conditioning modulates the reconstitution of peripheral blood cells, but not hHSCs or hMPPs in the BM (Figures 1D and 1E). These cellular and clonal-level analyses demonstrate that conditioning does not influence hHSC and hMPP engraftment, but affects the magnitude of hHSPCs’ hematopoietic contribution and alters the composition and timing of their hematopoietic output, indicating complex effects on both HSPCs and their niche. Mouse HSC clones exhibit similar changes in clonal expansion and lineage bias in response to irradiation conditioning, as previously reported ^12,23^. These findings could guide the clinical selection of transplantation regimens to strategically promote the reconstitution of specific immune cell types most critical for improving patient outcomes in a given clinical context.

Lastly, our results illustrate the limited efficiency of human hematopoiesis in xenograft mouse models by tracking human hematopoiesis at the clonal level through serial transplantation. Xenograft models are widely used in preclinical research to enable *in vivo* studies ^26,76,77^. However, the extent to which these models recapitulate human hematopoietic dynamics remains incompletely understood, particularly in the context of secondary transplantation. Our clonal analyses provide important data delineating the blood reconstitution of hHSCs and hMPPs in xenograft mouse models. We show that only a limited number of clones contribute to hematopoietic regeneration in each recipient mouse (Figures 2A and S3A), indicating that a small subset of transplanted hHSPCs dominates hematopoietic output. In secondary recipients, we found a marked reduction in overall human hematopoietic reconstitution in the peripheral blood (Figures 1F and 1H), with particularly low contributions to myeloid cells and T cells (Figure 1G). Unexpectedly, clonal tracking revealed that more hHSPC clones contributed to blood cells in secondary recipients compared to primary recipients (Figure 4), suggesting that some previously quiescent or latent hHSPC clones were activated upon serial transplantation, potentially in response to the inefficient hematopoiesis. Furthermore, we found that the majority of hHSPC clones contributing to BM hematopoiesis also contributed to EMH, a hallmark of hematopoietic stress (Figures 5F-5H). Altogether, these findings underscore the inadequate support of human hematopoiesis in xenograft mouse models.

Our study has some limitations in data collection. First, to preserve sufficient numbers of donor cells for transplantation into multiple secondary recipients, we could not analyze clonal tracking barcodes in the BM cells from the primary recipients. Second, we recovered clonal tracking barcodes from genomic DNA and were unable to analyze anucleated cells, such as red blood cells and platelets. Third, although the NSGW41 mouse model is widely used for studying human hematopoiesis and in preclinical research, it is known to provide limited support for human myelopoiesis and erythropoiesis. Despite these limitations, this study offers valuable insights into human hematopoiesis and the impact of pre-transplant conditioning.

## Conclusion

In conclusion, our findings reveal functional distinctions between hHSCs and hMPPs, elucidate the specific influences of pre-transplant conditioning regimens on human hematopoietic reconstitution, and illustrate the strengths and limitations of xenograft mouse models. Through quantitative in vivo clonal tracking and multi-tissue analyses, this study advances understanding of human hematopoiesis and provides a framework for optimizing experimental models, improving transplantation strategies, and informing the development of HSC-based cell and gene therapies.

## Abbreviations

hHSCs: Human hematopoietic stem cells
hMPPs: Human multipotent progenitors
hHSPCs: Human hematopoietic stem and progenitor cells
nEMH: Non-splenic extramedullary hematopoiesis
MNC: Mononuclear cells
BM: Bone marrow
hWBC: Human white blood cells
B: B cells
T: T cells
GR: Granulocytes
MO: Monocytes

## Declarations

### Data Availability

Data supporting the findings of this study and code used for data analysis and figure generation are available from the corresponding author upon request.

### Author Contributions

M.V.S. and R.L. conceived and designed the experiments. M.V.S. performed the experiments. P.C. and J.E. developed custom Python code for data analysis. M.V.S., P.C., J.E., and R.L. analyzed the data. M.V.S. and R.L. wrote the manuscript. All authors reviewed and approved the manuscript.

## Acknowledgements

We thank Dr. A. Nogalska and S. Lydon for assistance with manuscript editing; J. Contreras and B. Masinsin for FACS core management; and D. Ruble from the CHLA Genomics Core for high-throughput sequencing support. We are also grateful to all members of the Lu laboratory for their valuable discussions and insights.

