## Supplementary file for "Tracking Human HSCs and MPPs in Mice Reveals Distinct Clonal Dynamics and Responses to Pre-transplant Conditioning"

### Supplementary Table 1-2 and Supplementary Figure 1-5

### Supplementary Table 1. Transplantation setting for primary recipients

| Donor cells | Conditioning regimen | Number of mice |
|---|---|---|
| HSC: 7,000<br>MPP:18,000 | Busulfan (20mg/kg) | 5 |
| | Irradiation (100 cGy) | 5 |
| | Unconditioned | 5 |

### Supplementary Table 2. Transplantation setting for secondary recipients

| Primary mice | Conditioning regimen for 1° and 2° recipients | Number of secondary recipients | Donor cells for secondary recipients |
|---|---|---|---|
| B.1 | Busulfan | 4 | HSC: 3,000<br>MPP:130,000 |
| B.2 | Busulfan | 4 | HSC: 3,000<br>MPP:120,000 |
| B.3 | Busulfan | 4 | HSC: 2,500<br>MPP:125,000 |
| B.4 | Busulfan | 5 | HSC: 3,500<br>MPP: 130,000 |
| I.1 | Irradiation | 3 | HSC: 2,000<br>MPP: 110,000 |
| I.2 | Irradiation | 3 | HSC: 2,000<br>MPP: 115,000 |
| U.1 | Unconditioned | 4 | HSC: 2,500<br>MPP: 120,000 |
| U.2 | Unconditioned | 3 | HSC: 2,500<br>MPP: 110,000 |
| U.3 | Unconditioned | 4 | HSC: 2,500<br>MPP: 110,000 |

### Figure S1.

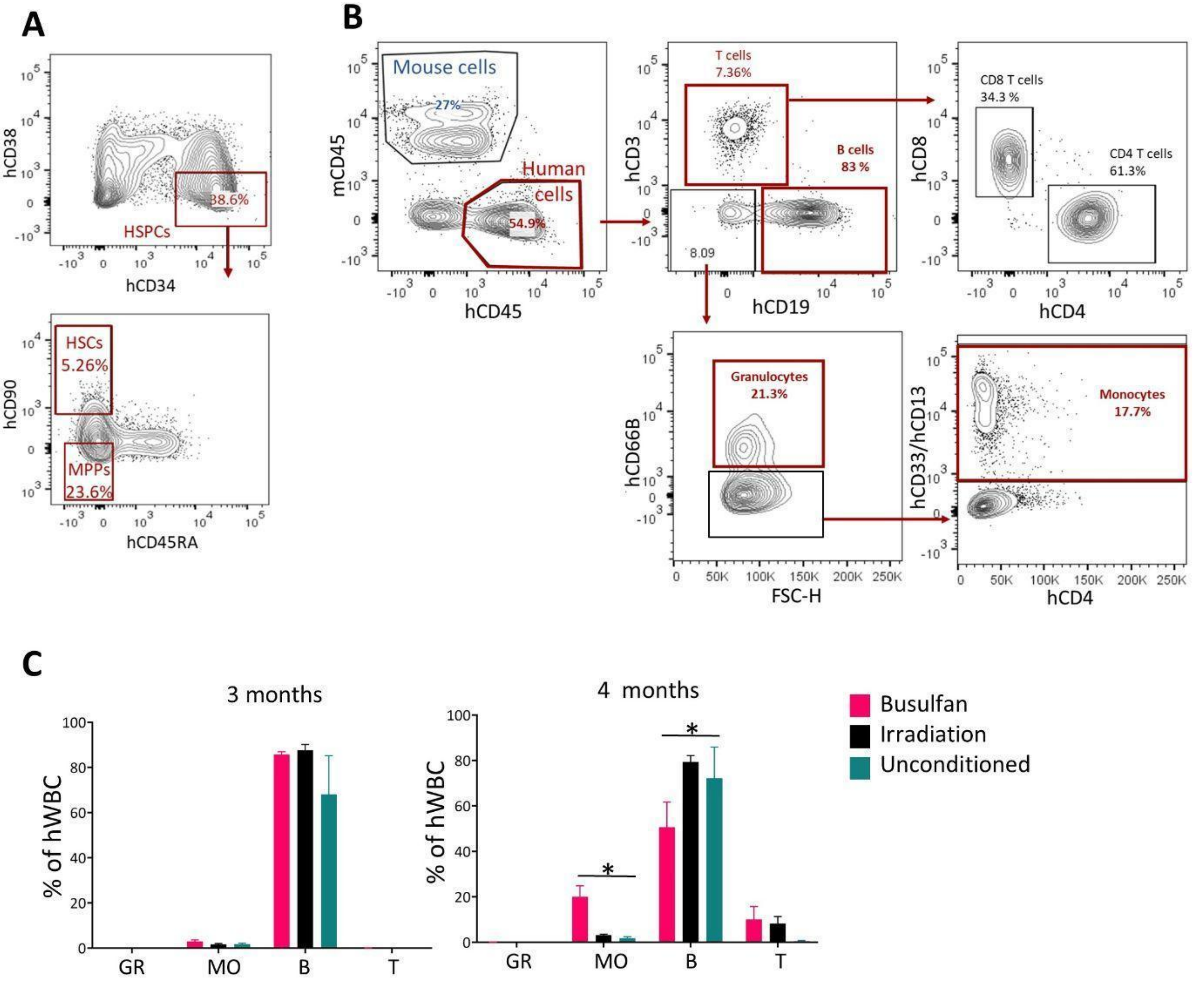


**Supplementary Figure 1. Tracking hHSPC differentiation in xenografted mice under various pre-transplant conditioning. (A)** Flow cytometry gating strategy for isolating donor hHSCs and hMPPs from umbilical cord blood. **(B)** Flow cytometry gating strategy for isolating hHSPC-derived blood cells from xenografted mice. **(C)** Composition of human-derived white blood cells (hWBCs) in the peripheral blood of primary recipients at 3 and 4 months post-transplantation**.** GR, granulocytes; MO, monocytes; B, B cells; T, T cells. Student's t-test; *$p<0.05$.

**Figure S2.**

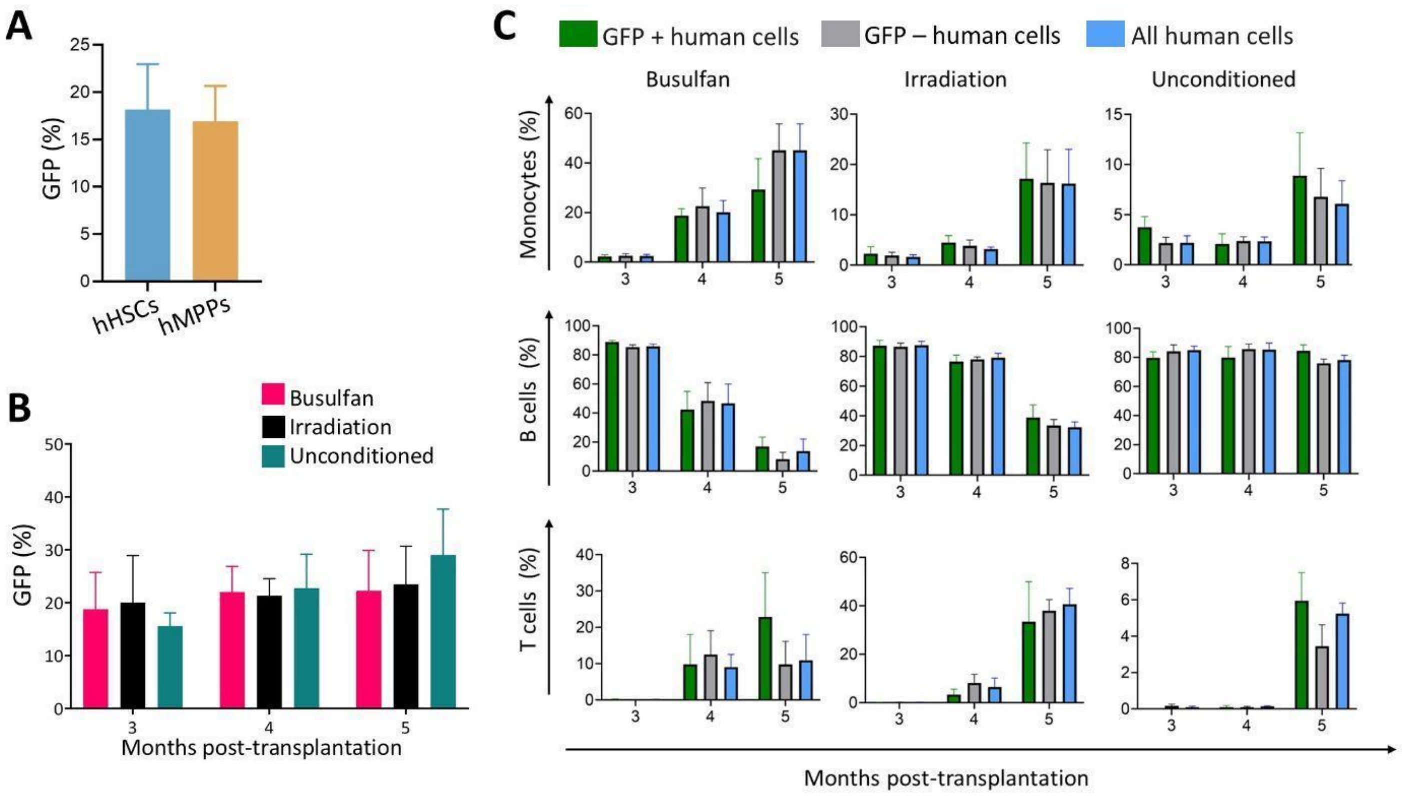


**Supplementary Figure 2. Comparison of barcoded and non-barcoded human hematopoietic cells.** Lentivirally transduced cells are barcoded and express GFP. **(A)** Transduction efficiency of hHSCs and hMPP, measured at 72 hours post-transduction. Data are presented as mean + SEM. n=6 using three different viral libraries. **(B)** Transduction efficiency of hHSPCs, measured by their hematopoietic progeny ($hCD45^{+}$) in the peripheral blood of recipient mice. **(C)** Cell type composition of barcoded ($GFP^{+}$), non-barcoded ($GFP^{-}$), and all human hematopoietic cells in the peripheral white blood cells from primary recipient mice over time.

**Figure S3.**

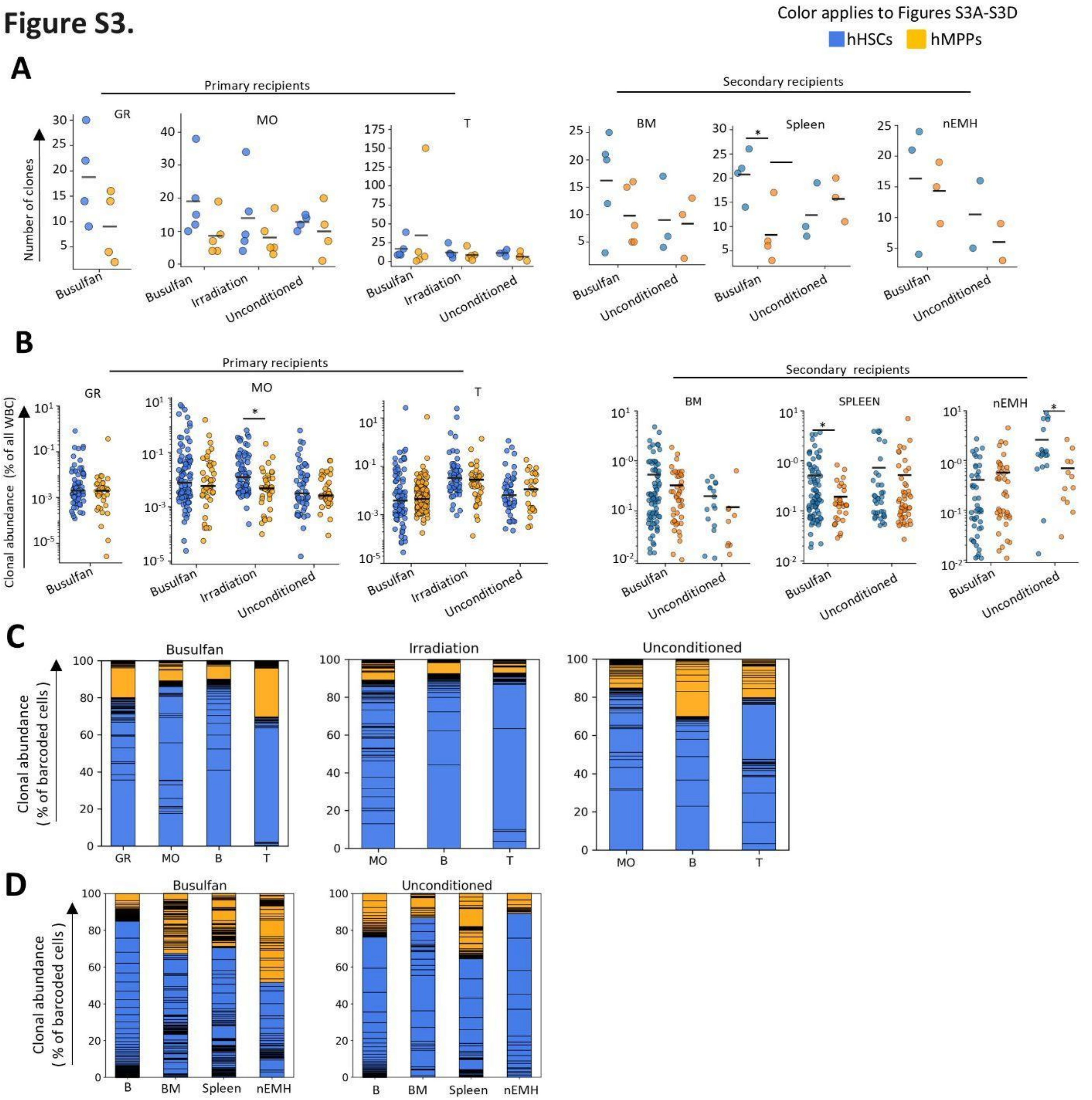


**Supplementary Figure 3. Contribution of hHSCs and hMPPs to various hematopoietic cell types and tissues.** Data was collected from the primary recipients 5 months post-transplantation and from the secondary recipients 12 months after secondary transplantation. **(A)** Number of hHSC and hMPP clones contributing to various hematopoietic cell types and tissues. Each dot represents data from one recipient mouse. Black bars indicate the means. Student's t-test; $*p<0.05$. **(B)** Clonal contributions of hHSCs and hMPPs to various hematopoietic cell types and tissues. Each dot represents a unique clone. Black bars indicate the medians. Mann–Whitney test; *p<0.05. **(C)** Relative contributions of hHSCs and hMPP clones to various hematopoietic cell types in the primary recipients. Each section within the columns represents one clone. Clones are ordered by their B cell contribution. **(D)** Relative contributions of hHSC and hMPP clones in the secondary recipients. Each section within the columns represents one clone. Clones are ordered by their B cell contribution. Gr, granulocytes; MO, monocytes; B, B cells; T, T cells; nEMH, non-splenic extramedullary hematopoietic sites.

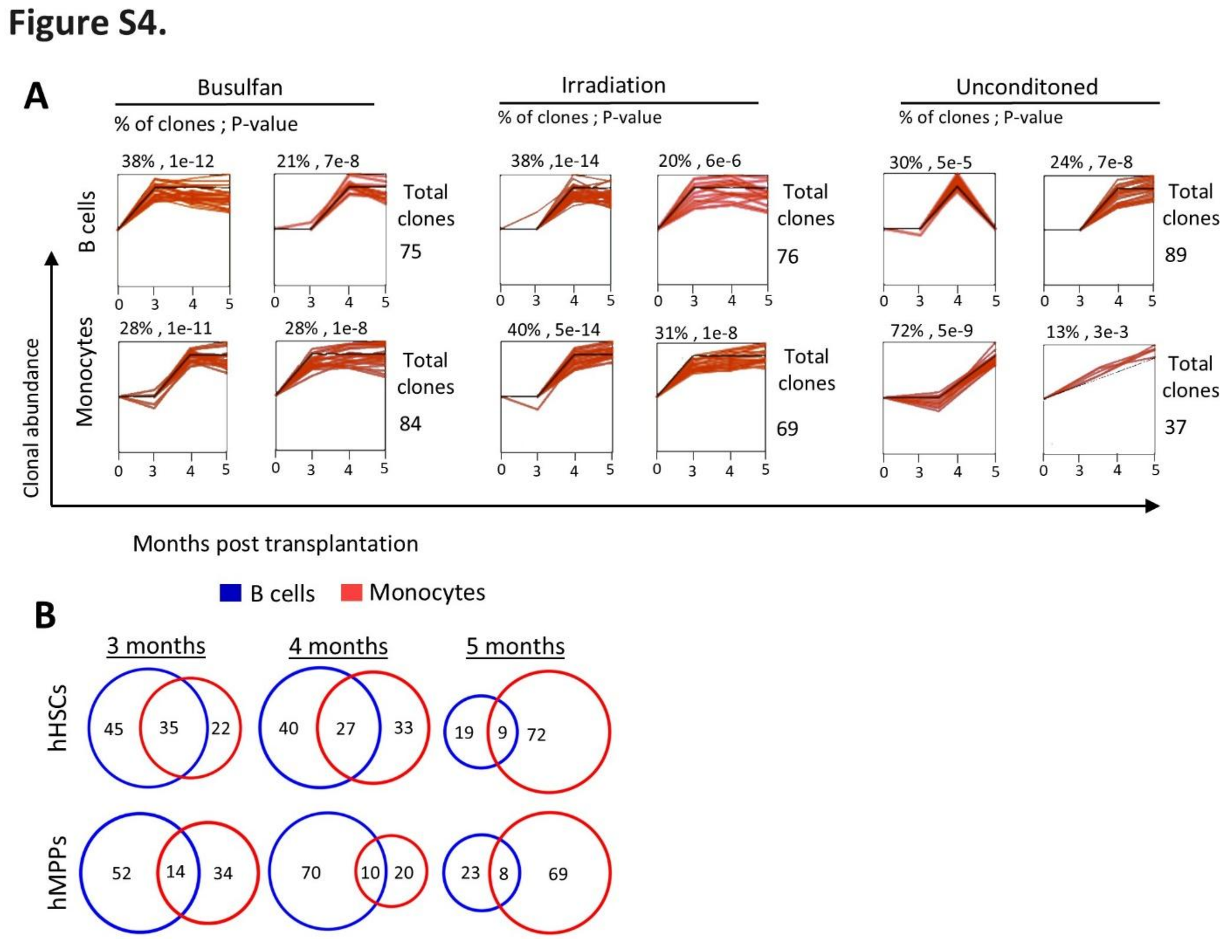


**Supplementary Figure 4. Temporal dynamics of hHSPC-derived blood cell regeneration in primary recipients following various pre-transplant conditioning regimens. (A)** Short Time-series Expression Miner (STEM[56]) analysis showing temporal clustering of clonal contributions from hHSCs and hMPPs to B cells and granulocytes in peripheral blood. Each panel depicts a cluster of clones with similar temporal dynamics. Shown are the two most prevalent temporal patterns within each group. Numbers above each panel denote the percentage of total clones in the cluster and the corresponding p-value relative to the expected proportion. Each red line represents a single hHSPC clone, and the black line shows the average temporal pattern for the cluster. Clones derived from hHSPCs and hMPPs were combined in this analysis. **(B)** Percentage of hHSC and hMPP clones that began contributing to B cells, monocytes, or both at various time points post-transplantation.

### Figure S5.

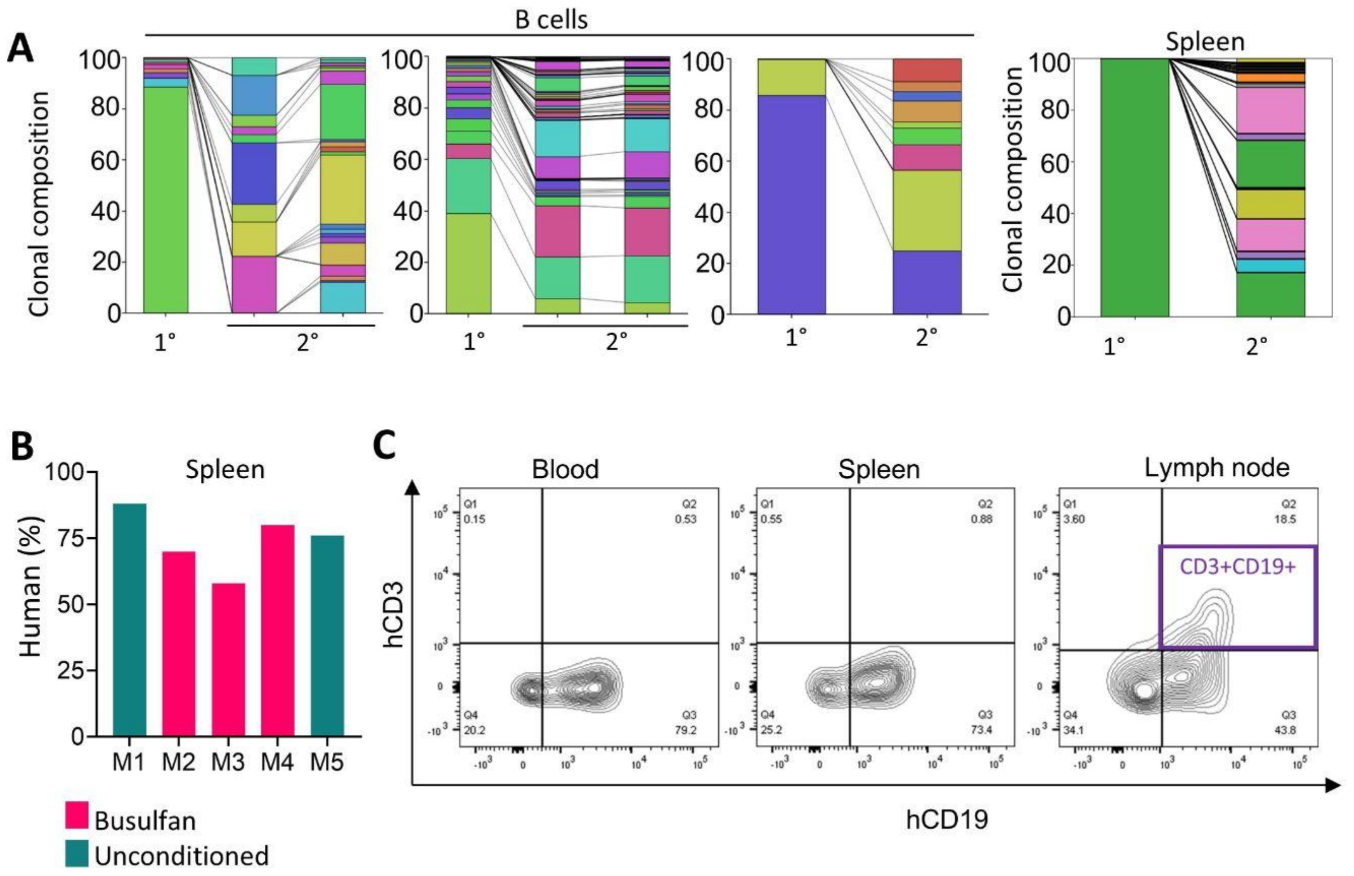


**Supplementary Figure 5. Comparing clonal contributions of hHSPCs between primary and secondary recipients. (A)** Clonal composition of human B cells in the peripheral blood (left) and human hematopoietic cells (hCD45+) in the spleen (right), comparing a primary recipient (1°) with its derived secondary recipients (2°). Each color represents a unique clone. Each column shows one mouse. **(B)** Percentage of human hematopoietic cells ($hCD45^{+}$) in the spleen. **(C)** Representative flow cytometry analysis (M4) showing the presence of $CD19^{+}CD3^{+}$ cells exclusively in the lymph node.
